# Discovery of genes that modulate flavivirus replication in an interferon-dependent manner

**DOI:** 10.1101/2021.07.20.453077

**Authors:** Sarah Lesage, Maxime Chazal, Guillaume Beauclair, Damien Batalie, Elodie Couderc, Aurianne Lescure, Elaine Del Nery, Frédéric Tangy, Annette Martin, Nicolas Manel, Nolwenn Jouvenet

**Author notes:** These authors contributed equally (Sarah Lesage, Maxime Chazal).

## Abstract

Establishment of the interferon (IFN)-mediated antiviral state provides a crucial initial line of defense against viral infection. Numerous genes that contribute to this antiviral state remain to be identified. Using a loss-of-function strategy, we screened an original library of 1156 siRNAs targeting 386 individual curated human genes in stimulated microglial cells infected with Zika virus (ZIKV), an emerging RNA virus that belongs to the flavivirus genus. The screen recovered twenty-one potential host proteins that modulate ZIKV replication in an IFN-dependent manner, including the previously known IFITM3 and LY6E. Further characterization contributed to delineate the spectrum of action of these genes towards other pathogenic RNA viruses, including Hepatitis C virus and SARS-CoV-2. Our data revealed that APOL3 acts as a proviral factor for ZIKV and several other related and unrelated RNA viruses. In addition, we showed that MTA2, a chromatin remodeling factor, possesses potent flavivirus-specific antiviral functions. Our work identified previously unrecognized genes that modulate the replication of RNA viruses in an IFN-dependent way, opening new perspectives to target weakness points in the life cycle of these viruses.

## Introduction

Viruses are high on the list of global public health concerns, as illustrated by recent epidemics caused by Ebola, Zika (ZIKV) and Nipah viruses, as well as by the ongoing SARS-CoV-2 pandemic. The vast majority of these emerging RNA viruses have zoonotic origins and have recently crossed host species barrier [1]. In order to establish itself in a host species, one of the first and most restrictive barriers that a virus needs to overcome is the antiviral innate immune system. This response has evolved to rapidly control viral replication and limit virus spread *via* detection of viral nucleic acids by pathogen recognition receptors (PRRs) [2]. These PRRs can be membrane-associated, such as Toll-like receptor (TLRs), or cytosolic, such as retinoic acid inducible gene I (RIG-I)-like receptor (RLRs). Upon binding to viral nucleic acids, these PRRs interact with adaptor proteins and recruit signaling complexes. These events lead to the expression of type I interferons (IFNs). Secreted IFNs will then bind to their heterodimeric receptor (IFNAR1/IFNAR2) and activate the canonical JAK/STAT pathway in infected and surrounding cells. This activation triggers the assembly of the interferon-stimulated gene 3 (ISGF3) complex (composed of STAT1, STAT2 and IRF-9 proteins), which subsequently induces the expression of up to approximately 2000 IFN-stimulated genes (ISGs) [3, 4], effectively establishing the antiviral state. ISGs comprise a core of genes that are induced at high levels essentially in all cell types, as well as cell-type specific genes that are the result of transcriptome remodeling [5, 6], highlighting the importance of studying ISGs in relevant cell types. Some of these ISGs have been well characterized. They directly block the viral life cycle by targeting specific stages of virus replication, including entry into host cells, protein translation, replication or assembly of new viral particles [3, 7]. Some ISGs are specific to a virus or a viral family, while others are broad-spectrum. They can also be negative or positive regulators of IFN signaling and thus facilitate, or not, the return to cellular homeostasis. However, the contribution of most ISGs to the antiviral state remains poorly understood.

Over the last decades, flaviviruses have provided some of the most important examples of emerging or resurging diseases, including ZIKV, dengue virus (DENV), Yellow fever virus (YFV) and West Nile virus (WNV) [8]. These flaviviruses are arthropod-borne viruses transmitted to vertebrate hosts by mosquitoes. They cause a spectrum of potentially severe diseases including hepatitis, vascular shock syndrome, encephalitis, acute flaccid paralysis, congenital abnormalities and fetal death [8]. They are now globally distributed and infect up to 400 million people annually. Lesser-known flaviviruses are beginning to emerge in different parts of the world, as illustrated by the recent incursion of Usutu virus (USUV) in the Mediterranean basin [9].

ZIKV was isolated in 1947 in a macaque from the Zika Forest in Uganda [10]. For decades, it remained in Africa and Asia where it sparked local epidemics characterized by a mild self-limiting disease in humans. In recent years, Asian lineage viruses have emerged as a global public health threat with widespread epidemics in the Pacific Islands and Americas, where over 35 countries have reported local transmission in 2016. An estimated 1 million individuals were affected by ZIKV in Brazil in 2015-16. Infection by ZIKV has been linked to several neurological disorders, including Guillain-Barré syndrome (GBS), meningoencephalitis, myelitis and congenital microcephaly, fetal demise and abortion [10]. Children exposed to ZIKV in utero may present neurocognitive deficits, regardless of head size at birth. ZIKV infection is now identified as a sexually-transmitted illness as well [11]. As all flaviviruses, ZIKV is an enveloped virus containing a positive-stranded RNA genome of ~11 kb. Upon viral entry, the viral genome is released and translated by the host cell machinery into a large polyprotein precursor. The latter is processed by host and viral proteases into three structural proteins, including C (core), prM (precursor of the M protein) and E (envelope) glycoproteins, and seven non-structural proteins (NS) called NS1, NS2A, NS2B, NS3, NS4A, NS4B and NS5 [8]. The structural proteins constitute the viral particle, while NS proteins coordinate RNA replication, viral assembly and modulate innate immune responses.

The importance of IFN signaling in mediating host restriction of ZIKV is illustrated by the severe pathogenicity in IFNAR1-/- and STAT2-/- but not in immunocompetent mice [12–14]. Moreover, the Zika strain that is responsible for the recent epidemics has accumulated mutations that increase neurovirulence via the ability to evade the immune response [15]. Microglial cells, which are the resident macrophages of the brain, represent ZIKV targets and potential reservoirs for viral persistence [16]. Moreover, they may play a role in ZIKV transmission from mother to fetal brain [17] and affect the proliferation and differentiation of neuronal progenitor cells [18]. In order to comprehend the molecular bases behind the efficacy of the IFN response to ZIKV replication, we set up a high throughput assay to identify genes that are modulating viral replication in human microglial cells (HMC3) stimulated with IFN.

## Results

### A loss of function screen identified genes modulating ZIKV replication in IFN-stimulated human microglial cells

We first performed pilot experiments to determine the feasibility of conducting large-scale loss-of-function studies to identify novel genes regulating ZIKV replication in stimulated HMC3 cells. Five hundreds cells were seeded in 384-well microplates on day 1, transfected with individual siRNA 6 hours later, treated with IFNα2 at day 2, infected with ZIKV at day 3 and fixed 24 hours later (Fig. 1A). Percentages of infected cells were determined by confocal analysis by measuring the number of cells expressing the viral E protein, using the pan-flavivirus anti-E antibody 4G2 (Fig. 1A). Nuclei were identified with DAPI staining for imaging and segmentation purposes. We optimized IFNα2 concentration and viral multiplicity of infection (MOI) to obtain a significant decrease of E-positive cells upon IFNα2-treatment (Fig. S1A). We used siRNAs targeting IFNAR1, which are expected to neutralize IFN signaling, as positive controls (Fig. S1A). siRNAs against IFITM3, an ISG known to potently inhibit ZIKV replication in several human cell lines and primary fibroblasts [19, 20], were used as additional positive controls (Fig. S1A). Negative controls were non-targeting siRNAs. As expected, in the presence of siRNAs targeting IFNAR1, IFN signaling was neutralized and the level of infection was almost rescued to the level of non-treated cells (Fig. S1A). In cells silenced for IFITM3 expression, the number of infected cells was partly restored to the level of non-treated cells (Fig. S1A). Such partial rescue was expected since the antiviral state requires the concerted action of numerous ISGs [21]. These data also revealed that IFITM3 is a potent anti-ZIKV ISG in microglial cells.

**Figure 1.**
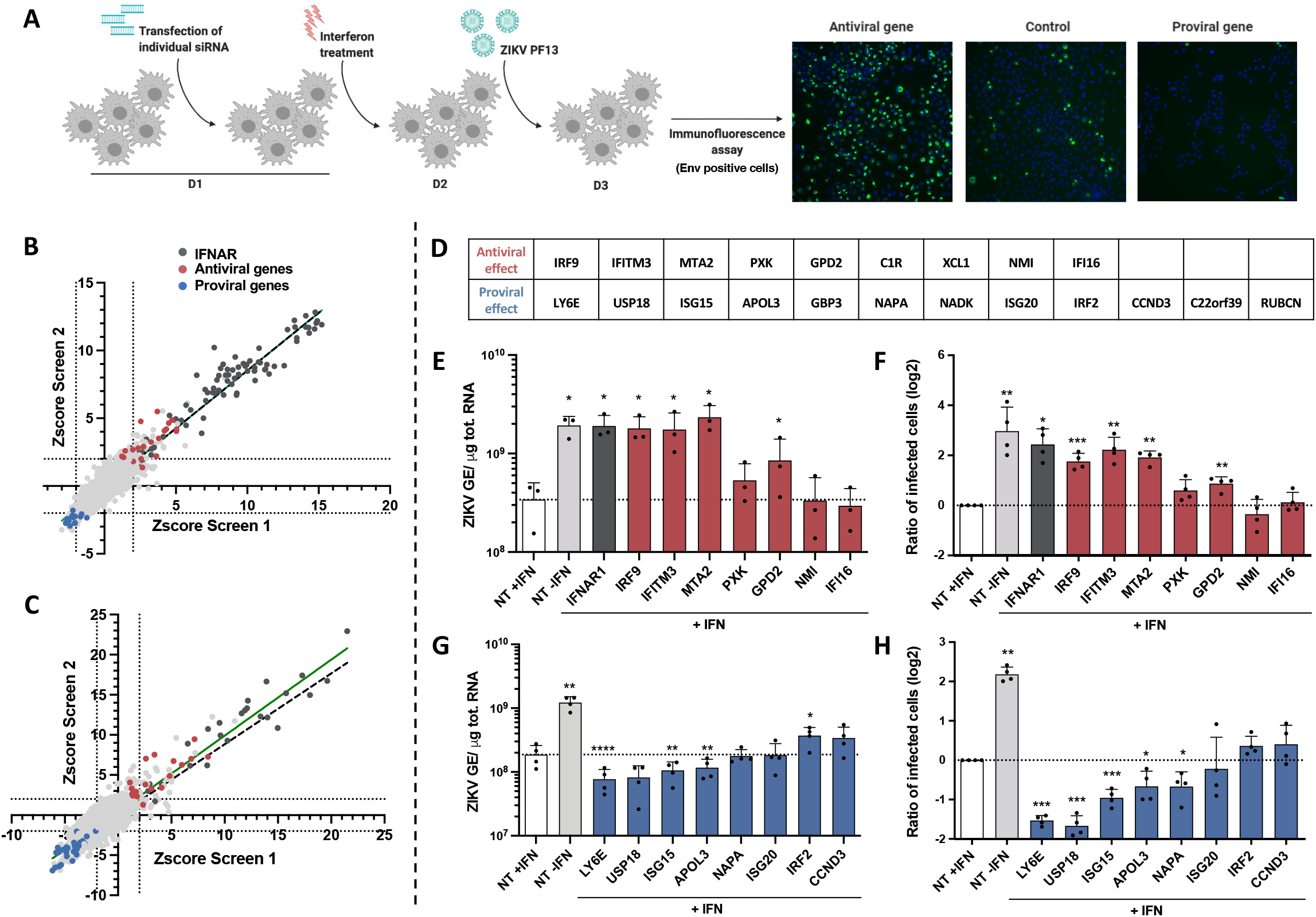
A loss of function screen identified genes modulating ZIKV replication in IFN-stimulated human microglial cells. (A) Scheme summarizing the screen conditions. (B, C) Scatter plots showing the rZ-score obtained in the 1st (B) and second analysis (C) of the 2 screens. The green line represents the linear regression, as compared to the expected perfect correlation (dotted black line). Antiviral and proviral hits are depicted in red and blue, respectively. (D) List of the antiviral and proviral hits as identified by the 2 analysis of the 2 screens. Assessment of the antiviral (E, F) and proviral (G, H) activities of some hits. HMC3 cells were transfected with either pool of 3 siRNAs against the indicated candidate gene or non-targeting (NT) siRNAs, treated with IFNα2 (200U/mL) for 24 hours and infected with ZIKV (at an MOI of 5 PFU/cell) for 24 hours. Control cells transfected with NT siRNA in the absence of IFNα2 treatment (NT-IFN) are included. (E, G) Cell-associated viral RNA was quantified by RT-qPCR and expressed as genome equivalents (GE) per μg of total cellular RNA. (F, H) The number of cells positive for viral protein E was assessed by flow cytometry and are expressed relatively to the NT+IFN control of each experiment. Data are means ± SD of three or four independent experiments, *p<0.05, **p<0.01,***p<0.001, paired t-tests.

We scaled up the experiment by screening an arrayed library containing 1158 siRNAs targeting 386 human genes (Table S1). These genes were selected based on a gene signature defined by clustering and correlation of expression with MX1, a well-described ISG, in a dataset of gene expression in primary human CD4^+^ T cells (Cerboni et al., in preparation). 36% of the identified genes overlapped with previous ISG libraries [21, 22], ensuring that the screen would be simultaneously capable of identifying expected positive hits and find new genes of interest. To overcome potential off-target effect and a limited efficacy of transcript knockdown, each gene was targeted by 3 different siRNAs. Numerous siRNAs targeting IFNAR1 and IFNAR2 were used as positive controls. Negative controls were non-targeting siRNAs. Transfection efficiency was evaluated using siRNAs against KIF11, a protein essential for cell survival [23]. The same experimental protocol than in small-scale experiments was applied (Fig. 1A). Three images were acquired per condition with an INCell2200 automated wide-field system. The mean cell count and the percentages of infected cells were extracted from quantification. The screen was performed in duplicate.

For quality control purposes, we first compared the number of cells in each well in the 2 replicates. We observed an expected distribution of the number of cells in 3 fields with a median close to 1000 cells per well for the two replicates (Fig. S1B). The number of cells per condition was slightly higher in the first replicate than in the second one. However, the R^2^ coefficient of determination of the linear regression was close to 0.7 (Fig. S1C), indicating that the reproducibility of the experiment was correct. As expected [23], siRNAs against KIF11 were lethal, validating the transfection protocol (Fig. S1B, C). The 2 screens were first analyzed by taking into consideration the intensity of the E signal per cell. The number of cells expressing the viral protein E distributed as predicted, with a median close to 15% for the 2 screens (Fig. S1D). As expected from pilot experiments (Fig. S1A), siRNAs targeting IFNAR-1 and −2 rescued ZIKV replication in IFN-treated cells (Fig. S1D, E). The reproducibility of the infection status of the cells between the 2 screens, with a R2 greater than 0.8, was satisfactory (Fig. S1E). The data were then analysis using a second approach that identified infected cells based on the E expression independently of the intensity of the signal. The 2 methods identified similar number of infected cells (Fig. S1F). Results were expressed as robust Z-scores for each siRNA (Fig. 1B, C). Genes were defined as hits when at least two over three of their robust Z scores had an absolute value superior to 2 in the two replicates, in at least one of the analysis. The screen identified 9 antiviral genes and 12 proviral ones (Fig. 1D). Some hits were previously described as modulators of ZIKV replication, such as IFITM3 [19, 20] and LY6E [24], thus validating our loss-of-function screening approach. These twenty-one hits were selected for further validation.

IFNα2-treated HMC3 cells were transfected with pool of 3 siRNAs against each candidate, and not by individual ones as in the primary screening. Twenty-four hours post-ZIKV infection, intracellular viral RNA production was quantified by RT-qPCR and the number of cells positive for the viral protein E was assessed by flow cytometry analysis. The same samples were used to assess the efficacy of the siRNAs. RT-qPCR analyses revealed that 15 out of the 21 siRNA pools were reducing the expression of their respective targets in IFN-treated cells (Fig. S1G). mRNAs levels of C1R, XCL1, GBP3, NADK, C22orf39 and RUBCN were below the detection limit in IFNα2-treated HMC3 (Fig. S1G). These genes were thus excluded from further anaysis. Reduced expression of IRF9, IFITM3, MTA2 and GPD2 significantly enhanced both viral RNA yield and the number of infected cells as compared to IFNα2-treated cells transfected with control siRNAs (Fig. 1E and F). Both IRF9, which belongs to the ISGF3 complex [25], and IFITM3 [26] are well-known broadly-acting IFN effectors. The activities of MTA2 have, so far, not been linked to antiviral immunity. MTA2 is a component of the nucleosome remodeling deacetylase NuRD complex, which exhibits ATP-dependent chromatin remodeling activity in addition to histone deacetylase activity [27]. Ten times more viral RNA copies were recovered in cells silenced for MTA2 expression than in control cells (Fig. 1E) and four times more cells were positive for the viral E protein (Fig. 1F). These effects were comparable to the ones induced by IFNAR1 silencing (Fig. 1E and F). Transfection with siRNA against the other 3 antiviral candidates (PXK, NMI and IFI16) had no significant effect on ZIKV replication in these assays (Fig. 1E and 1F), suggesting that they may be false positive candidates. Reducing the expression of LY6E, ISG15 and APOL3 significantly decreased both viral RNA production and the number of cells positive for the E protein (Fig. 1G and 1H), validating the pro-viral activities of these 3 candidates. The pro-viral function of USP18 and NAPA were also validated since reducing their expression led to a significant reduction of the number of infected cells as compared to control cells (Fig. 1H). Reduced expression of ISG20 or CCND3 had no significant effect on ZIKV replication (Fig. 1G and 1H). IRF2, which was identified as a pro-viral hit by the screen, behaved like an antiviral gene in the validation experiments (Fig. 1G). Together, these experiments validated the antiviral function of IRF9, IFITM3, MTA2 and GPD2 and the pro-viral function of LY6E, USP18, ISG15, APOL3 and NAPA in IFNα2-treated HMC3 cells infected with ZIKV.

### Effect of a selection of candidate genes on HCV and SARS-CoV-2 replication

We next explored the ability of 10 candidate genes (IRF9, IFITM3, MTA2, GPD2, LY6E, USP18, ISG15, APOL3, GBP3 and NAPA) to modulate the replication of two other pathogenic RNA viruses: Hepatitis C virus (HCV) and Severe Acute Respiratory Syndrome Coronavirus 2 (SARS-CoV-2), which are, respectively, related and unrelated to ZIKV. HCV, which is a member of the Hepacivirus genus within the *Flaviviridae* family, has a tropism for hepatocytes. SARS-CoV-2 belongs to the *Coronaviridae* family and has a tropism for pneumocytes and enterocytes.

HCV infections were conducted in hepatoma Huh-7.5 cells, which support well viral replication [28]. Huh-7.5 cells are unable to induce IFN expression since they express an inactive form of RIG-I [29] but they possess an intact JAK/STAT pathway and do thus respond to IFN treatment [30]. RT-qPCR analyses revealed that 8 out 10 siRNA pools efficiently reduced the expression of their respective targets in stimulated Huh-7.5 cells (Fig. S2A). Since LY6E and APOL3 mRNA levels were under the limit of detection of the assays in IFNα2-treated Huh-7.5 cells (Fig. S2A), they were excluded from further analysis. As expected, reduced expression of IFNAR and IRF9 significantly enhanced HCV RNA yield and the production of infectious particles in IFNα2-treated Huh-7.5 cells, as compared to control cells (Fig. 2A, B). Reduced expression of IFITM3, MTA2 or GPD2, which significantly enhanced ZIKV replication in HCM3 cells (Fig. 1E, F), did not affect HCV RNA production (Fig. 2A). However, surprisingly, their reduced expression triggered a significant decrease in the release of infectious HCV particles as compared to control cells (Fig. 2B). This suggests that they might favor a late stage of HCV replication cycle. RT-qPCR analysis and titration assays identified USP18 and ISG15 as pro-HCV factors in IFNα2-treated Huh-7.5 cells (Fig. 2C, D), validating previous results [31–34]. Of note, HCV RNA production and infectious particle release were significantly increased in cells with reduced NAPA levels (Fig. 2C, D), suggesting that NAPA may exert an antiviral effect on HCV, while it was not the case for ZIKV (Fig. 1G, H).

**Figure 2.**
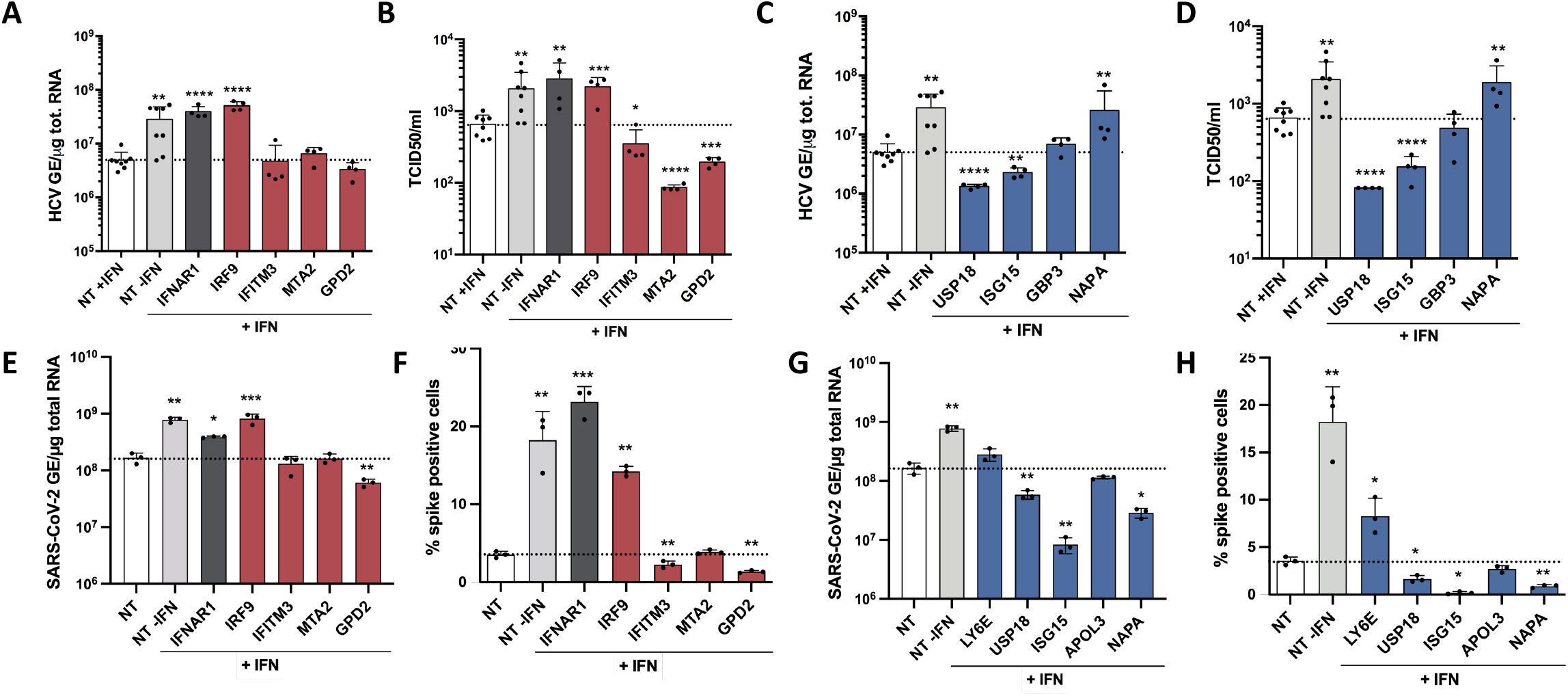
Effect of a selection of candidate genes on HCV and SARS-CoV-2 replication. (A-D). Huh-7.5 cells were transfected with a pool of 3 siRNAs against selected candidates (antiviral and proviral genes, as identified in the ZIKV screen (Fig. 1), are in red and blue, respectively) or non-targeting (NT) siRNAs, treated with IFNα2 (200U/mL) for 24 hours, then infected with HCV at a MOI of 3 TCID50/cell for 48 hours. (A, C) Cell-associated viral RNA was quantified by RT-qPCR and expressed as genome equivalents (GE) per μg of total cellular RNA. (B, D) Release of infectious HCV particles was determined by TCID50 assays. Data are expressed relatively to the NT+IFN control of each experiment. Plotted values are expressed relative to mean NT+IF across experiements and represent means ± SD of two independent experiments each in duplicates, *p<0.05, **p<0.01,***p<0.001, ****p<0.0001 paired t-tests. (E-H). A549-ACE2 cells were transfected with a pool of 3 siRNAs against selected candidates (antiviral genes in red, proviral genes in blue) or non-targeting (NT) siRNAs, treated with IFNα2 (200U/mL) for 24 hours and infected with SARS-CoV-2 at a MOI of 2 for 24 hours. (E, G) Cell-associated viral RNA was quantified by RT-qPCR and expressed as genome equivalents (GE) per μg of total cellular RNA. (F, H) The number of cells positive for the viral protein spike (S) was assessed by flow cytometry Data are expressed relatively to the NT+IFN control of each experiment. Data are means ± SD of triplicates of one experiment, *p<0.05, **p<0.01,***p<0.001, paired t-tests.

SARS-CoV-2 replication was assessed in A549 alveolar epithelial cells expressing the viral receptor ACE2 (A549-ACE2) by RT-qPCR and flow cytometry analysis using an antibody against the viral protein Spike (S). Of note, silencing GBP3 in A549-ACE2 cells triggered cell death. RT-qPCR analyses showed that all siRNA pools were reducing the expression of their respective targets in stimulated A549-ACE2 cells (Fig. S2B). These analyses revealed the ability of IRF9 to act as an anti-SARS-CoV-2 gene (Fig. 2E, F). Unexpectedly, GPD2 and IFITM3, which we identified as genes possessing anti-ZIKV activities (Fig. 1), tended to behave like pro-viral genes in the context of SARS-CoV-2 infection (Fig. 2E and 2F). Viral RNA yields decreased significantly in cells silenced for USP18, ISG15 and NAPA expression, as compared to cells transfected with control siRNAs (Fig. 2G), suggesting that these 3 genes promote viral replication in stimulated A549-ACE2 cells. By contrast to what we observed in ZIKV infected cells (Fig. 1G, H), LY6E seemed to restrict SARS-CoV-2 (Fig. 2H). These results are in accordance with a recent report [35]. Reducing MTA2 or APOL3 expression did not affect SARS-CoV-2 replication.

These results suggest that some genes are broadly-acting IFN effectors, such as IRF9 and ISG15. Other genes appeared to have evolve modulatory function toward a specific viral family or genus, such as APOL3 and MTA2. Finally, some genes, including LY6E, IFITM3 and GPD2, exhibited opposite modulatory functions towards different viral species.

### ZIKV, DENV-2, WNV, VSV and MeV, but not MVA, require the expression of APOL3 for optimal replication in IFN-treated cells

LY6E, ISG15 and APOL3 exhibited significant pro-ZIKV activities in stimulated cells, as measured by cell-associated viral RNA levels (Fig. 1G) and percentage of E-positive cells (Fig. 1H). Among these 3 genes, APOL3 is the least described and was thus selected for further characterization. APOL3 is one of the 6 members of the apolipoprotein L gene family. Apolipoproteins are typically associated with the transport of lipids in the organism and were originally described as members of the high-density lipoprotein family, which are involved in cholesterol transport [36]. In human cells, the expression of the 6 members of the APOL gene family are up-regulated by multiple pro-inflammatory signaling molecules, including IFNs and TNFα [36, 37]. These regulations suggest a link between APOL proteins and the innate immune system. siRNA targeting APOL2, APOL3, APOL4, APOL5 and APOL6 were present in our library (Table S1). Among these 5 APOLs, only APOL3 was identified as a facilitator of ZIKV infection by our screen (Fig. 1). We decided to test the ability of APOL1 to modulate ZIKV replication since it was previously identified in a high-throughput overexpression screen as an ISG able to increase YFV infection in *STAT1*^-/-^ fibroblasts and Huh-7cells [21].

Analysis of mRNA levels of APOL1 and APOL3 revealed that the genes were upregulated by IFNα2 treatment in HMC3 cells (Fig. 3A). Both genes thus qualify as genuine ISGs in these cells. The implication of APOL1 and APOL3 in ZIKV replication was investigated using loss-of-function approaches. siRNA-silencing reduced the levels of APOL1 and APOL3 mRNAs by ~80% and ~85%, respectively, when compared to cells expressing scrambled control siRNAs (Fig. 3B). USP18, which is known to negatively regulates the JAK-STAT pathway, and, as such, is a broad-spectrum pro-viral factor [38], was identified during our screen as a pro-ZIKV candidate in HMC3 cells (Fig. 1D). Since its pro-ZIKV function was validated in our system (Fig. 1H), siRNAs specific for USP18 were used as positive controls. siRNA-silencing reduced the abundance of USP18 mRNAs by ~80% when compared to cells expressing control siRNAs (Fig. 3B). Viral replication was assessed by flow cytometry by measuring the number of cells positive for the viral protein E in cells silenced for APOL1, APOL3 or USP18, treated or not with IFNα2. Since ZIKV is sensitive to IFNα2-treatment (Fig. S1A), higher MOIs were used in IFNα2-treated cells than in untreated ones to compensate for its antiviral effects. As expected (Fig. 1H), reduced expression of USP18 significantly decreased the number of IFN-treated cells positive for the viral protein E, as compared to cells transfected with control siRNAs (Fig. 3C). Extinction of APOL1 and APOL3 resulted in a modest, but reproducible, decrease in the number of E-positive HMC3 cells pre-treated with IFNα2 (Fig. 3C). A pro-viral effect of APOL1 was also observed in unstimulated cells (Fig. 3C). The efficacy of the siRNAs against APOL1 and USP18 were further validated by Western Blot analysis using specific antibodies (Fig. 3D). APOL3 levels in cell lysates could not be assessed due to the lack of available antibodies. Levels of expression of the viral proteins NS5 and E were slightly decreased in IFNα2-cells expressing reduced levels of APOL1 or APOL3, compared to control cells (Fig 3D). Together, these results suggest that ZIKV requires the expression of APOL1 and APOL3 for optimal replication in HMC3 cells. By contrast to APOL1, the pro-viral action of APOL3 was dependent on IFNα2-treatment.

**Figure 3.**
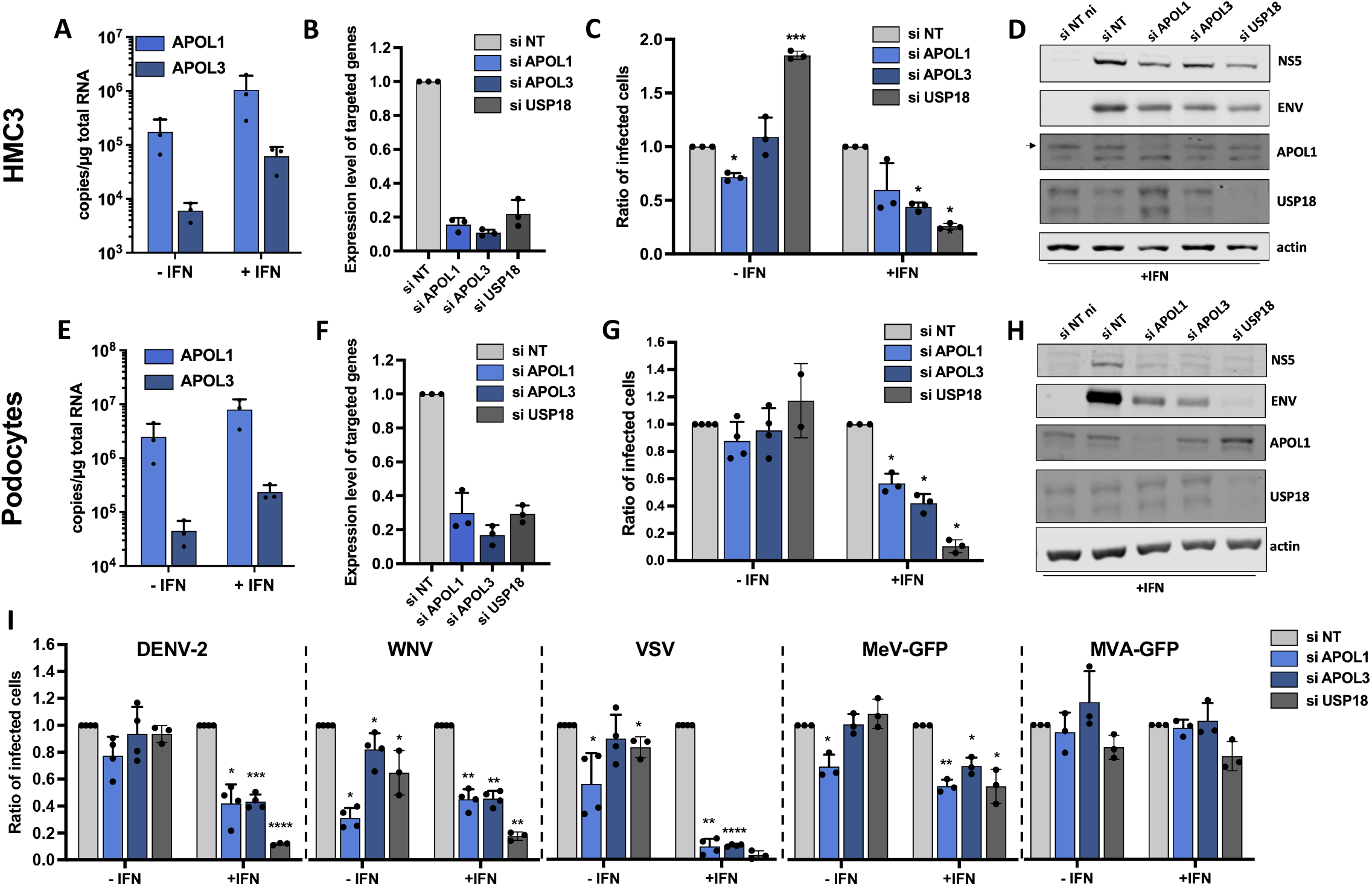
Effect of reduced expression of APOL1 and APOL3 on the replication of a panel of viruses in HMC3 cells and podocytes. (A, E) *APOL1* mRNA and *APOL3* mRNA abundance were quantified by RT-qPCR analysis in HMC3 cells or podocytes treated or not with IFNα2 (200U/mL) for 24 hours and expressed as copy numbers per μg of total cellular RNA. HMC3 cells (B) or podocytes (F) were transfected with pool of 3 siRNAs targeting *APOL1, APOL3* and *USP18* mRNAs or with non-targeting (NT) control siRNAs. The relative amounts of *APOL1, APOL3* and *USP18* mRNAs were determined by RT-qPCR analysis and were normalized to that of GAPDH mRNA. They are expressed as compared to abundance relative to cells transfected with control NT siRNAs. HMC3 cells (C) or podocytes (G) were transfected with the indicated siRNAs, treated, or not, with IFNa2 (200U/mL) for 24 hours, and infected with ZIKV for 24 hours. HMC3 cells were infected at a MOI of 2 and podocytes at a MOI of 1. The percentages of cells that were positive for the viral E proteins were determined by flow cytometric analysis. Data are expressed relatively to the siRNA NT control of each experiment. HMC3 cells (D) or podocytes (H) were treated with IFNα2 (200U/mL), transfected with the indicated siRNAs pools and subjected to Western blotting analysis with antibodies against the indicated proteins. (I) HMC3 cells were transfected with the indicated siRNAs pools, treated with IFNa2 (200U/mL) for 24 hours and infected with the indicated viruses for 18 to 24 hours, at the MOI indicated in the MM section. The percentages of the cells positive for viral proteins or GFP were determined by flow cytometric analysis. Data are means ± SD of three or four independent experiments, *p<0.05, **p<0.01,***p<0.001, paired t-tests.

To ensure that the APOL1- and APOL3-mediated modulation of viral replication was not restricted to HMC3 cells, silencing experiments were performed in ZIKV-infected human podocytes treated or not with IFNα2. Podocytes are physiologically relevant for ZIKV infection since viral RNA was detected in kidneys of infected patients [39]. Assessing the mRNA abundance of APOL1 and APOL3 by RT-qPCR analysis of cells treated or not with IFNα2 revealed that both genes qualify as ISGs in podocytes (Fig. 3E). siRNA-mediated silencing of APOL1, APOL3 and USP18 was efficient in podocytes (Fig. 3F). Reduced expression of APOL1 or APOL3 resulted in a significant decrease of the percentage of infected cells in IFNα2-treated podocytes, but not in unstimulated cells (Fig. 3G). Western blot analysis performed in IFNα2-treated podocytes revealed that cells expressing little APOL1/3 were producing less viral proteins than controls cells (Fig. 3H), confirming the pro-ZIKV activity of the two APOLs. These data revealed that APOL3 and APOL1 facilitate the replication of ZIKV in podocytes treated with IFNα2.

We tested whether APOL1 and APOL3 were active against DENV-2 or WNV, which are mosquito-borne flaviviruses closely related to ZIKV. HMC3 cells were treated or not with IFNα2 and the MOIs were adapted to the IFNα2 treatment. Flow cytometry analysis using anti-E antibodies revealed that both DENV-2 and WNV replication were significantly decreased in IFNα2-treated cells silenced for APOL1 or APOL3 expression (Fig. 3I). Reducing APOL1 and APOL3 expression in non-treated cells also significantly reduced WNV replication (Fig. 3I). Thus, APOL1/3 may well have flavivirus genus-specific proviral activities since they seems to contribute to ZIKV, WNV and DENV replication (Fig. 3C, D, G, H and I) but not to SARS-CoV-2 replication (Fig. 2G, H). To further delineate the spectrum of action of these two genes towards other viruses, we tested the effect of APOL1/3 silencing on the replication of Vesicular Stomatitis virus (VSV) and Measles virus (MeV), which are negative-strand RNA viruses belonging to the *Rhabdoviridae* and *Paramyxoviridae* families, respectively. Experiments were performed with a MeV strain modified to express GFP [40]. We also included in the analysis Modified Vaccinia Ankara virus (MVA), a DNA virus belonging to the *poxviridae* family, that was engineered to express GFP (MVA-GFP). Flow cytometry analysis using an antibody against the viral protein G revealed that VSV was highly dependent on APOL1 and APOL3 expression for efficient replication in IFNα2-treated HMC3 cells (Fig. 3I). Optimal replication of MeV-GFP in stimulated HMC3 cells also required APOL1 and APOL3 expression (Fig. 3I). APOL1 proviral activity was also observed in unstimulated cells (Fig. 3I). By contrast, MVA-GFP replication was not affected by reduced expression of either APOL1 or APOL3 (Fig. 3I).

Together, these experiments suggest that APOL1 and APOL3 could favor a replication process shared by ZIKV, DENV-2, WNV, VSV and MeV. Unlike APOL1 in HMC3 cells, APOL3 pro-viral activities were dependent on IFN-treatment.

### APOL1 and APOL3 likely promote viral replication independently of their interaction with phosphoinositides

Recent data revealed that APOL1 and APOL3 play a role in lipid metabolism in podocytes and, more specifically, in the regulation of the production of phosphatidylinositol-4-phosphate (PI(4)P), via an indirect interaction with the PI4KB kinase [41]. PI(4)P is involved in Golgi secretory functions by facilitating the recruitment of proteins that promote vesicular transport [42]. PI(4)P is also essential for the establishment of efficient viral replication via the formation of membranes which serve as platforms for the production of viral RNA [43, 44]. APOL1 and/or APOL3 could thus impact ZIKV replication via their ability to regulate the production of PI(4)P. To test this hypothesis, we first assessed APOL1 and APOL3 localization in HMC3 cells. In the absence of specific antibodies for APOL1 and APOL3 validated for immunofluorescence assays, we investigated the localization of GFP-tagged versions of APOL3 and APOL1 in HMC3 cells, together with markers for the Golgi apparatus (Fig. 4A), early or late endosomes (Fig. S3). APOL1-GFP and GFP-APOL3 localized in closed proximity to the cis-Golgi (Fig. 4A), where PI(4)P and PI4KB localize [45], and not in late nor early endosomes (Fig. S3). In line with this, APOL1-GFP and GFP-APOL3 associated with PI4KB in HMC3 cells (Fig. 4B). Of note, APOL1-GFP was also detected in vesicle-like structures whose identity could not be established (Fig. 4A, white arrow). They may represent lipid droplets or fragmented Golgi. The localization of APOL1-GFP and GFP-APOL3 could not be investigated in ZIKV-infected cells since we observed that transfection rendered cells non-permissive to viral infection.

**Figure 4.**
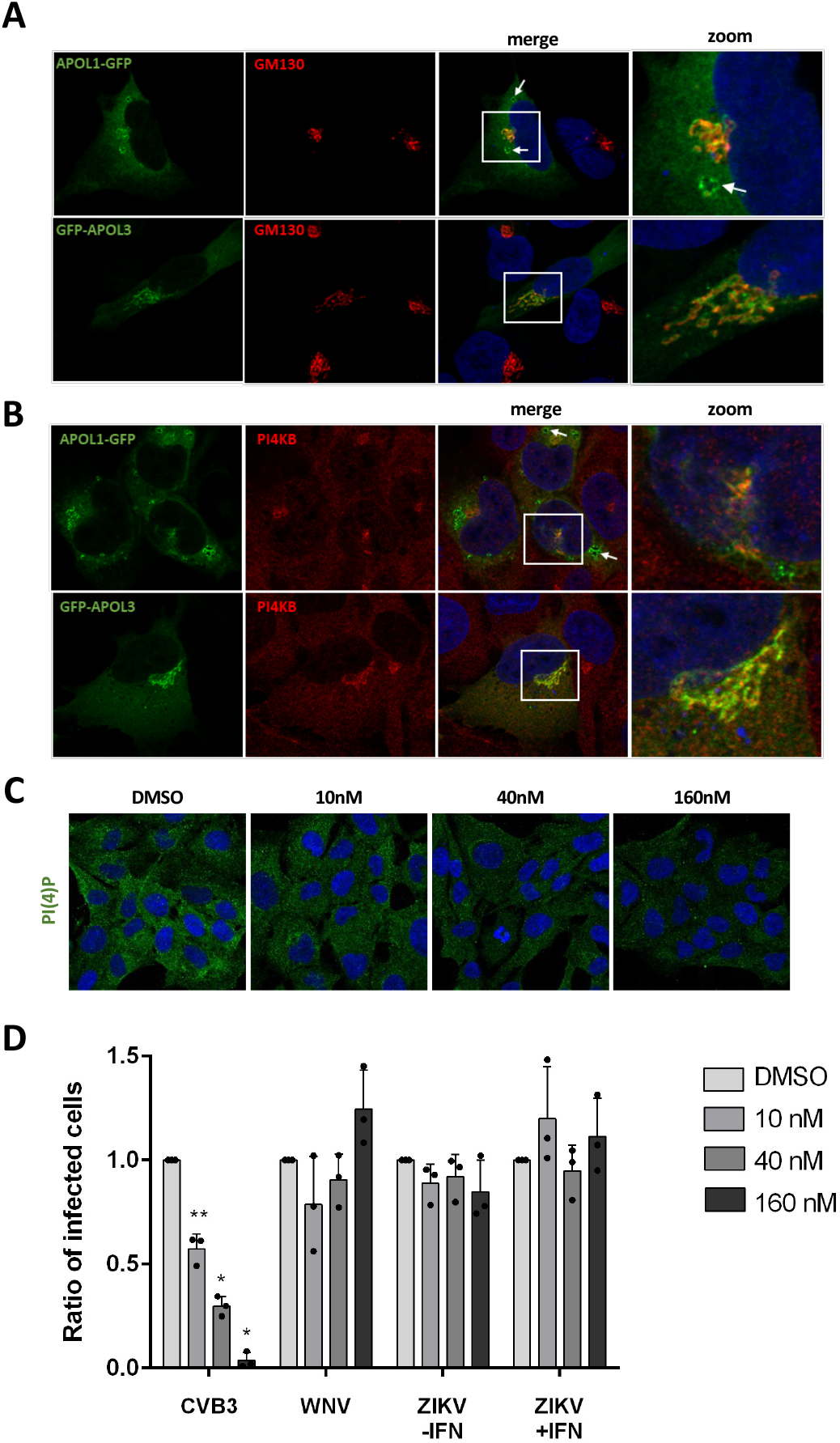
APOL1 and APOL3 promote viral replication independently of their interaction with phosphoinositides. HMC3 cells were transfected with GFP-tagged versions of APOL1 and APOL3. Thirty hours later, they were stained with antibodies recognizing GM130 (A) or PI4KB (B) and with NucBlue to detect nuclei. Images are representative of numerous observations over 2 independent experiments. The white arrow shows an APOL1-GFP-positive vesicle. (C) HCM3 cells were treated with different concentrations of the PI4KB inhibitor and were stained for PI(4)P. (D) HMC3 cells treated with different doses of PI4KB inhibitor were infected with CVB3, WNV or ZIKV, in the presence or absence of IFNα2 (200U/mL). The percentages of the cells positive for viral proteins were determined by flow cytometric analysis. Data are means ± SD of three independent experiments, *p<0.05, **p<0.01,***p<0.001, one-way ANOVA.

We then performed experiments with a well-characterized PI4KB kinase inhibitor that decreases PI(4)P expression [46]. We first analyzed by immunofluorescence the intensity of the PI(4)P signal in cells treated for 24 h with different concentrations of the drug in HCM3 cells. The presence of the PI4KB inhibitor triggered a dose-dependent decrease of the PI(4)P signal (Fig. 4C), suggesting that the drug is efficient in HMC3 cells. We then infected cells with ZIKV in the presence of different concentration of the inhibitor. Since the effect of APOL3 on ZIKV replication is dependent on IFNα2 (Fig. 3), the analysis was also performed in stimulated cells. Coxsackie B3 virus (CVB3), an enterovirus that replicates in a PI(4)P-dependent manner, was used as a positive control since its replication is sensitive to the drug [47]. As negative controls, we used cells infected with WNV, whose replication is not affected by the PI4KB inhibitor [48]. As previously shown in HeLa cells [47], a dose-dependent reduction of the number of cells positive for the CVB3 viral protein 1 (VP1) was triggered by the inhibitor treatment (Fig. 4D). As shown previously in monkey cells [48], WNV replication was unaffected by the PI4KB inhibitor in HCM3 cells (Fig. 4D). ZIKV protein production was not sensitive to the treatment with the PI4KB kinase inhibitor, independently of the presence of IFNα2 (Fig. 4D). These experiments suggest that the pro-viral activities of APOL1 and APOL3 are not related to their interaction with PI4KB or PI(4)P in microglial cells.

### MTA2 restricts ZIKV replication in IFNα2-stimulated human cells

MTA2 was identified in our screen as a gene with potent anti-ZIKV activities (Fig. 1). MTA2 shows a very broad expression pattern and is strongly expressed in many tissues. It belongs to the NuRD complex, which establishes transcriptional modulation of a number of target genes in vertebrates, invertebrates and fungi [27]. Since its function has, so far, not been linked to viral infection, we decided to investigate its potential antiviral activities further. To ensure that the MTA2-mediated inhibition of viral replication was not restricted to HMC3 cells, experiments were also performed in Huh-7 hepatoma cells, which are physiologically relevant for flavivirus infection and are thus extensively used in *Flaviviridae* research. siRNAs targeting IFNAR1 were used as positive controls in these experiments. siRNA-silencing reduced the levels of MTA2 and IFNAR1 mRNAs by at least 80%, when compared to cells expressing scrambled control siRNAs, independently of the stimulation or infection status of HMC3 and Huh-7 cells (Fig. 5A-D). MTA2 was included in our gene list because its expression clustered with MX1 in T cells (Cerboni et al. in preparation). However, MTA2 mRNA abundance, as measured by RT-qPCR analysis, remained unchanged upon IFNα2 treatment in both cells types (Fig. 5A and 5C), indicating that MTA2 is not an ISG in these cells.

**Figure 5.**
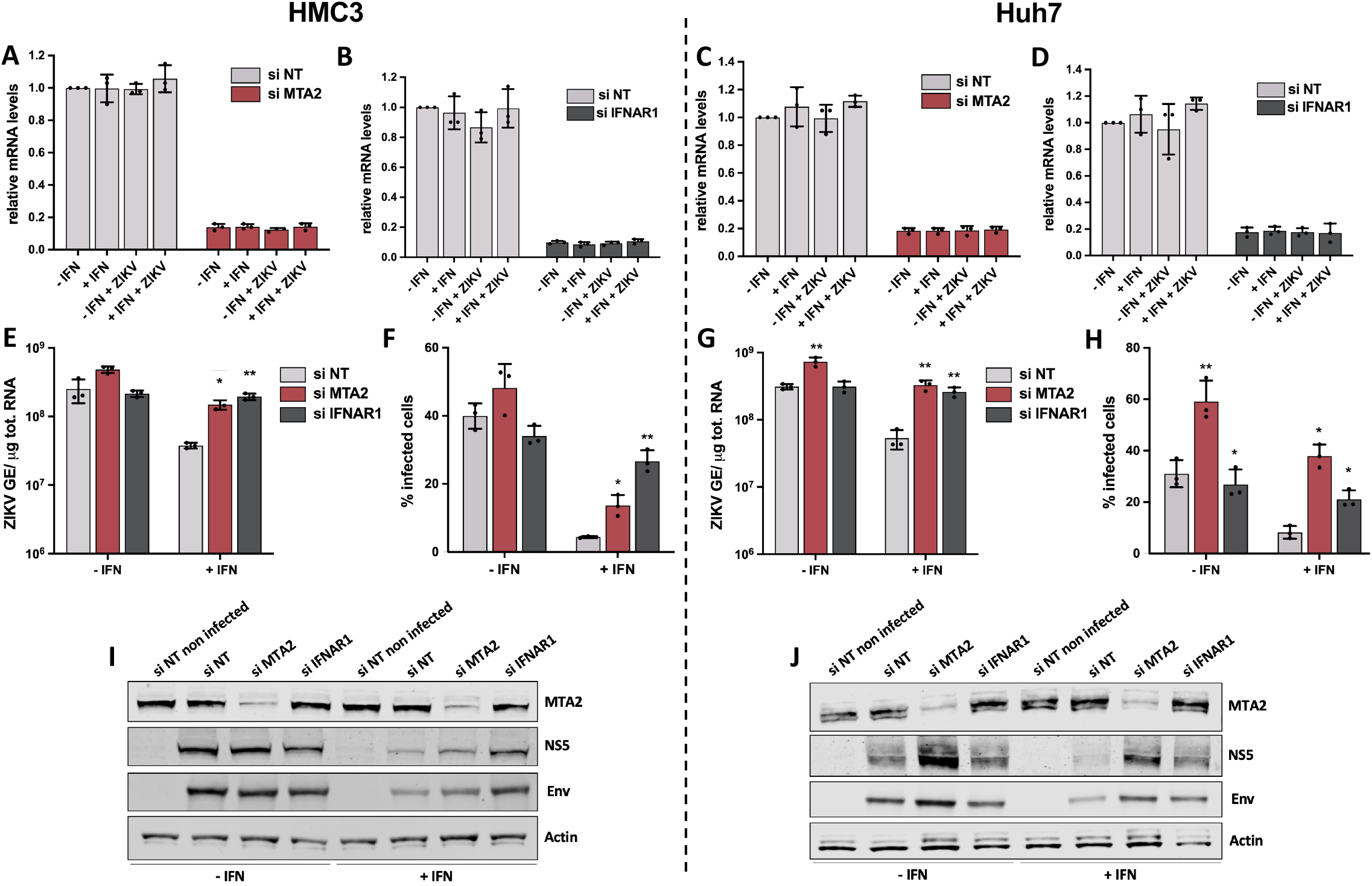
MTA2 restricts ZIKV replication in IFNα2-stimulated cells. HMC3 (A, B, E, F, I) and Huh-7 cells (C, D, G, H, J) were transfected with pool of 3 siRNAs targeting *MTA2* or *IFNAR1* mRNAs or non-targeting (NT) control siRNAs, treated or not with IFNα2 (100U/mL) for 24 hours, and infected with ZIKV (MOI of 1 for HMC3 cells, MOI of 5 for Huh-7 cells) for 24 hours. (A-D) The relative amounts of *MTA2* and *IFNAR1* mRNAs were determined by RT-qPCR analysis and normalized to that of *GAPDH* mRNA and siRNA-NT without IFN. (E, G) Cell-associated viral RNA was quantified by RT-qPCR and expressed as genome equivalents (GE) per μg of total cellular RNA. (F, H) Number of infected cells was assessed by staining of viral protein E and flow cytometry analysis. (I, J) Cells were treated with IFNα2 (200U/mL) or left untreated, transfected with the indicated siRNAs pools and subjected to Western blotting analysis with antibodies against the indicated proteins. Data are means ± SD of three independent experiments, *p<0.05, **p<0.01,***p<0.001, paired t-tests.

Assessment of viral replication by RT-qPCR revealed that cell-associated viral RNA yields were significantly higher in IFN-treated HMC3 cells silenced for MTA2 expression, as compared to controls cells (Fig. 5E). This is in line with previous results (Fig. 1E). Cytometry analysis using anti-E antibodies confirmed that MTA2 anti-ZIKV activities were dependent on the presence of IFN in HCM3 cells (Fig. 5F). Since MTA2 is not an ISG in HMC3 cells (Fig. 5A), these results suggest that MTA2 may require an active IFN signaling to exert its anti-ZIKV activities in these cells. As in HMC3 cells, reduced expression of MTA2 triggered a significant increase of intracellular viral RNA production in stimulated Huh-7 cells (Fig. 5G). Reducing MTA2 expression had a more pronounced effect on the percentage of infected cells that reducing IFNAR1 expression in stimulated Huh-7 cells (Fig. 5H). Albeit to a lesser extent than in stimulated cells, MTA2 anti-ZIKV activity was also observed in non-stimulated Huh-7 cells by flow cytometry and RT-qPCR analysis (Fig. 5G and H).

The effect of MTA2 on viral protein production was further assessed by Western blot analysis using anti-E and anti-NS5 antibodies in stimulated and unstimulated HMC3 and Huh-7 cells. These experiments validated further the efficacy of the siRNAs against MTA2 in both cell lines (Fig. 5I and 5J). Expression of the viral proteins NS5 and E were increased in stimulated HMC3 and Huh-7 cells expressing reduced levels of MTA2 or IFNAR1, as compared to control cells (Fig. 5I and 5J). In agreement with the flow cytometry analysis (Fig. 5H), MTA2 anti-ZIKV activity was less dependent of IFN-treatment in Huh-7 cells than in HMC3 cells (Fig. 5I and 5J). As observed in flow cytometry analysis (Fig. 5H), MTA2 effect on viral protein production was more potent than the one of IFNAR1 in stimulated Huh-7 cells (Fig. 5J).

These results represent the first evidence of the ability of MTA2 to restrict the replication of any virus.

### MTA2 restricts YFV and WNV replication in IFNα2-stimulated Huh-7 cells

We tested whether MTA2 was active against WNV and YFV in Huh-7 cells, which are permissive to these 2 flaviviruses. As in previous experiments, higher MOIs were used in the presence of IFNα2. Cytometry analysis revealed that MTA2 silencing significantly enhanced the replication of these 2 flaviviruses in an-IFN dependent manner (Fig. 6A and 6B), indicating that MTA2 antiviral activity is broader that ZIKV. We then tested the effect of MTA2 silencing on the replication of VSV and MeV in Huh-7 cells. Reduced expression of MTA2 decreased the number of cells infected with VSV and MeV (Fig. 6C and 6D), independently of the IFN stimulation. This is consistent with the pro-HCV activity of MTA2, as measured by titration in stimulated Huh-7.5 cells (Fig. 2B). MTA2 may thus possesses a flavivirus genus-specific antiviral function.

**Figure 6.**
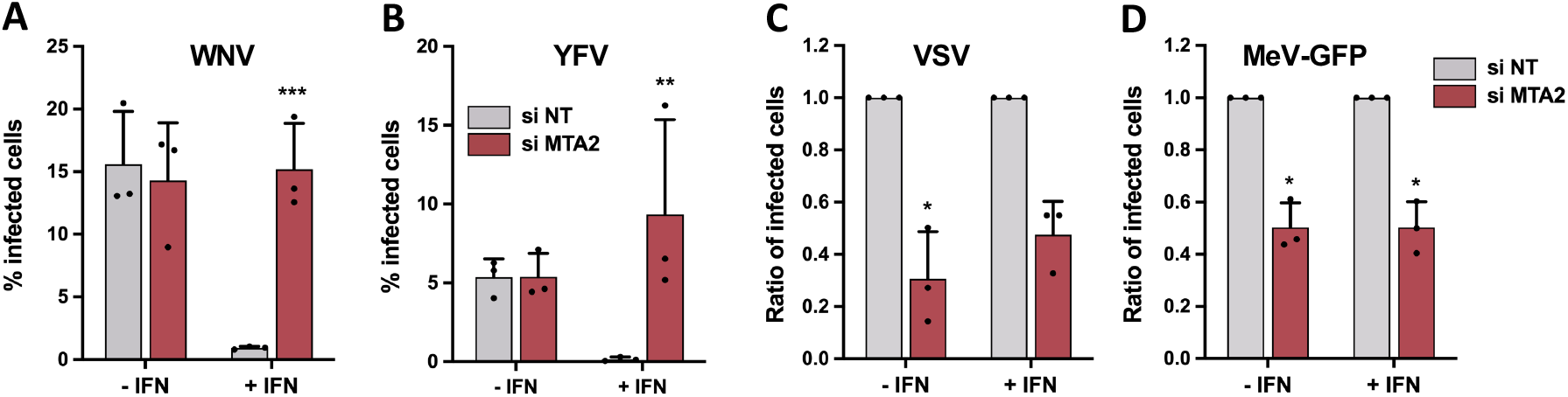
Effect of reduced expression of MTA2 on the replication of YFV, WNV, VSV and MeV-GFP. Huh-7 cells were transfected with the indicated siRNAs pool, treated or not with IFNα2 (200U/mL) for 24 hours and infected with WNV (A) or YFV (B) for 24 hours, at the MOIs indicted in the MM section. The percentages of the cells positive for viral protein Env was determined by flow cytometric analysis. HMC3 cells were transfected with the indicated siRNAs pool, treated or not with IFNα2 (200U/mL) for 24 hours and infected with VSV for 18 hours (C) or MeV-GFP for 24 hours (D), at the MOIs indicted in the MM section. The percentages of the cells positive for viral protein G or GFP were determined by flow cytometric analysis. Data are expressed relatively to the siRNA NT control of each experiment. Data are means ± SD of three or four independent experiments, *p<0.05, **p<0.01,***p<0.001, paired t-tests.

## Discussion

Several gain-of-fonction screens have been performed to identify ISGs that modulate flavivirus infection. Pionner screens tested the activities of relatively small amounts of ISGs by overexpression [49, 50]. The first comprehensive overexpression screen in which more than 380 ISGs were evaluated for antiviral activity against six viruses, including the *Flaviviridae* HCV, WNV and YFV, was published in 2011 by Schoggins and collaborators [21]. To avoid potential physiological irrelevance induced by gene overexpression, we opted for a silencing approach to identify genes that modulate ZIKV replication in an IFN-induced state. We used a siRNA library which was designed in the context of an HIV project. A limitation of our library is that targeted genes were selected based on a transcriptomic analysis of primary T cells stimulated by contacts with activated monocytes (Cerboni et al. in preparation), and not on ZIKV-target cells. Nevertheless, it contains a high fraction of core ISGs that overlaps with previous screens [21, 22]. Furthermore, most arrayed screens designed to identify cellular factors that modulate ZIKV replication, including ours, monitored viral replication after a single round of infection, often by assessing viral protein expression. Therefore, only genes that inhibit early stages of viral replication, up to protein production, can be identified. Quantifying viral titers in supernatants collected from individual wells of the first round of screening should identify genes that affect late stages of viral replication, such as viral assembly, maturation and release, as well as viral infectivity. Alternatively, viral replication could be monitored after several rounds of infection. Nevertheless, despite these two main limitations, our screening strategy identified 21 genes affecting the number of cells positive for the viral protein E in IFN-treated microglial cells.

Some hits were previously described as ISGs able to modulate ZIKV replication, such as IFITM3 [19, 20] and LY6E [24], thus validating our screening approach. Despite being in our gene list, Viperin, IFI6, PARP-12 and C19orf66, which are known to affect ZIKV replication in human cells [51–55], were not identified as viral modulators by our strategy. They may have a weaker influence on viral replication in HMC3 cells than in the cells in which their role was previously established [51–55]. In line with this hypothesis, Viperin restricts the replication of several neurotropic flaviviruses in a cell type-dependent manner [56]. One can also envisage that the expression levels of Viperin, IFI6, PARP-12 and C19orf66 are low in HMC3 cells, even when stimulated by IFNα2, and therefore are poorly, if at all downregulated by specific siRNAs.

We validated the role of 5 hits as genes contributing to an optimal ZIKV replication in stimulated HMC3 cells using RT-qPCR and flow cytometry analysis: LY6E, USP18, ISG15, APOL3 and NAPA. The identification of LY6E as a gene enhancing ZIKV replication was expected, since it was previously shown to promote the internalization of flaviviruses in U2OS human osteosarcoma cells [24]. Since ISG15 and USP18 negatively regulate IFN signaling pathway [57, 58], they are expected to act as broad pro-viral ISGs. NAPA interacts with SNARE protein complexes to trigger their disassembly [59]. SNARE proteins belong to a superfamily of membrane fusion proteins that localize at the plasma membrane, the Golgi apparatus and on different endocytic vesicles. They regulate the traffic of these vesicles between the plasma membrane and the Golgi [60]. Several viruses, including influenza A virus and VSV, hijack SNARE proteins to enter host cells [61]. SNARE complexes may thus contribute to NAPA pro-viral activities. We validated the anti-viral functions of 4 screen hits in stimulated HMC3 cells infected with ZIKV: IRF9, IFITM3, MTA2 and GPD2. Identification of IRF9, which plays a key role in ISG expression [25], and IFITM3, which restricts early stages of ZIKV infection [19, 20], validates our screening strategy. GPD2 is a mitochondrial protein involves in the metabolism of glycerol. No link between GPD2 and viral infections has been established yet. However, it regulates inflammatory response in macrophages [62]. Further experiments will be required to confirm that NAPA and GPD2 have the ability to modulate ZIKV replication in stimulated human cells.

Experiments performed on cells infected with HCV or SARS-CoV-2 contributed to delineate the spectrum of action of a selection of the screen hits. Our data illustrate once again that some ISGs have virus-specific antiviral activities [63]. For instance, we found that ZIKV, but not the related HCV, was sensitive to IFITM3 expression, confirming (HCV) and extending (ZIKV) recently reported data [20, 64]. Our data also confirm that some ISGs exert opposite effect on different viruses. For instance, as described previously, LYE6 promotes the replication of ZIKV [24] but restricts that of SARS-CoV-2 [35]. Viruses have developed numerous innovative strategies to evade ISG-mediated restriction [65, 66]. Hijacking individual ISG for promoting their replication is one of them. This hypothesis may explain why, to our surprise, our screen recovered more pro-viral genes that antiviral ones.

Our results identified APOL3 and APOL1 as ISGs required for optimal ZIKV replication in HMC3 cells. The proviral activity of APOL1 was less dependent on IFN that the one of APOL3, which suggest that both proteins act via different mechanisms. Reduced expression of APOL1 and APOL3 also restricted the replication of WNV and DENV in stimulated HMC3 cells. In line with these findings, over-expression of APOL1 was previously reported to increase YFV replication in Huh-7 cells [21]. VSV replication was highly reduced in the absence of one of these 2 genes. APOL1 has been reported to act as an antiviral ISG in the context of infection with alphaviruses (Sindbis virus and Venezuelan equine encephalitis virus) and human parainfluenza virus [21, 67]. Its over-expression also inhibits HIV-1 infection in monocytes [68]. Thus, APOL1 and APOL3 seem to behave like pro- or anti-viral ISG depending on the virus, or have no obvious role (HCV). In line with our data on flaviviruses, a recent report found that overexpression of APOL1 promoted infection with ZIKV and DENV-2, confirming a proviral role for this factor [69]. However in this study, an increase of ZIKV, DENV-2 and HCV replication was also reported in Huh-7.5 cells expressing siRNAs targeting APOL1 and APOL3 in the absence of IFN treatment [69], yielding conflicting data that will deserve further investigation.

From our data, we formulated the hypothesis that APOL1 and APOL3 pro-viral activities could be linked to their ability to bind to anionic phospholipids, including several phosphoinositides, in particular PI(4)P [41]. Both APOL1 and APOL3 were detected in PI(4)P-containing liposomes [41]. Moreover, reduced expression of APOL3 resulted in reduction of PI(4)P levels in podocytes [41]. PI(4)P plays a pivotal role in the Golgi secretory functions by facilitating recruitment of proteins that promote vesicular transport [42]. Our immunofluorescence data revealed that GFP-tagged version of APOL1 and APOL3 localize mainly in the Golgi, where PI(4)P localizes [42]. Numerous RNA viruses, including various *Picornaviridae*, HCV, coronaviruses and parainfluenza type 3, rely on PI(4)P to build membranous replication platform, where viral replication and assembly take place [70]. Flaviviruses are no exception and also trigger intracellular membrane remodelling for the building of membranous replication platforms [71]. Experiments conducted with a well-characterized PI4KB kinase inhibitor excluded the possibility that APOL1 and APOL3 pro-viral effects depend on PI(4)P and its synthetizing protein, PI4KB. In line with these results, PI(4)P are not important for the replication of WNV and the related Usutu virus [48]. Other avenues should be thus explored to understand how APOL1 and APOL3 modulate the replication of several RNA viruses. Since APOL1 and APOL3 impact the replication of unrelated viruses, they may be negative regulators of IFN signalling pathway, by, for instance, contributing to the proper routing of members of the JAK/STAT pathway.

Our data revealed that MTA2 possesses potent antiviral function in the context of ZIKV, WNV and YFV in stimulated cells, whereas its exhibited a proviral role for HCV, VSV and MeV. Reducing MTA2 expression in the presence of IFNα2 enhanced flaviviral replication to a level comparable to the inhibition of IFNAR1. Despite being part of our gene list, MTA2 is not induced by IFN in HMC3 Huh-7 cells but its antiviral activity was dependent on IFN in these cells. MTA2 may thus interact with an ISG to act on viral replication. MTA2 is a component of the NuRD complex, an unusual complex which exhibits ATP-dependent chromatin remodeling activity in addition to histone deacetylase activity [27]. The complex establishes transcriptional modulation of a number of target genes in vertebrates, invertebrates and fungi [27]. MTA2 related activities have not, so far, been linked to innate immunity in virus-infected cells. However, a link between the NuRD complex and STAT1-mediated IFN response was established in the context of infection with the protozoan parasite *Toxoplasma gondii* [72]. A *Toxoplasma* protein, named TgIST, translocates to the host cell nucleus where it recruits the complex NuRD to STAT1-dependent promoters, resulting in altered chromatin and blocked STAT1-mediated transcription [72]. Moreover, HDAC1, which is also a member of the NuRD complex, associates with both STAT1 and STAT2 in human cells [73]. Furthermore, specific reduction of HDAC1 expression inhibits IFNβ-induced transcription whereas HDAC1 overexpression enhances IFNβ-induced transcription [73]. Finally, HDAC inhibitors block the formation of ISGF3 and this was associated with impairment of STAT2 nuclear accumulation in mouse L929 cells [74]. These findings indicate a fundamental role for deacetylase activity and HDAC1 in transcriptional control in response to IFN. One could thus envisage that MTA2, within the NuRD complex, also interacts with STAT1 in cells stimulated with IFN and favors its action locally. This interaction could restrict flavivirus infection, possibly *via* enhancing the expression of a subset of flavivirus-specific ISGs.

Our work identified previously unrecognized genes that modulate the replication of RNA viruses in an IFN-dependent way. Future studies combining transcriptomic analysis of IFN-treated cells and high throughput loss-of-function screens will help define the interferome of cell types relevant for viral infection. Such studies are primordial to continue investigating the complexity the IFN-mediated antiviral program.

## Materials and Methods

### Cells

Human microglial cells (HMC3) were purchased from the American Type Culture Collection (ATCC, CRL-3304). They were maintained in Dulbecco’s modified Eagle’s medium (DMEM) containing GlutaMAX I and sodium pyruvate (Gibco), supplemented with 10% fetal bovine serum (FBS) and 1% penicillin-streptomycin (P/S) (final concentration of 100 units/mL and 100 μg/mL, respectively) (Sigma) and non-essential amino acids (GibcoTM NEAA 100X MEM, Life Technologies). Podocytes were described previously [75]. They were grown at 33°C in Roswell Park Memorial Institute medium (RPMI) containing GlutaMAX I (Gibco) and supplemented with 10% FBS and P/S. Before any experiments, cells were differentiated during 7 days at 37°C. Human hepatocellular carcinoma Huh-7 cells [76], which were kindly given by Cinzia Traboni (IRBM, Pomezia, Italy), were maintained in DMEM supplemented with 10% FBS and 1% P/S. Huh-7.5 cells (Apath, LLC), a subclone of Huh-7 cells [76] were cultured in DMEM supplemented with non-essential amino acids, 1mM sodium pyruvate, 10% FBS and P/S. Vero NK cells, which are African green monkey kidney epithelial cells, were purchased from ATCC and used for viral titration assays. They were maintained in DMEM containing GlutaMAX I and sodium pyruvate (Gibco), supplemented with 10% FBS and P/S. *Aedes albopictus* C6-36 cells were maintained in Leibovitz’s L-15 medium containing 10% FBS, 1% P/S, 1% Non-Essential Amino Acids Solution (Gibco) and 2% Tryptose Phosphate Browth (Gibco). Human lung epithelial A549 cells were modified to stably express hACE2 using the pLenti6-hACE2 lentiviral transduction, as described previously [77]. Cell cultures were verified to be mycoplasma free with the MycoAlertTM Mycoplasma Detection Kit (Lonza).

### Virus stocks, titration and infection

The Zika strain PF13 (kindly provided by V. M. Cao-Lormeau and D. Musso, Institut Louis Malardé, Tahiti Island, French Polynesia) was isolated from a viremic patient in French Polynesia in 2013. Stocks were produced on C6-36 cells. The Dengue 2 virus (DENV-2) strain Malaysia SB8553 was obtained from the Centro de Ingeniería Genética y Biotecnología (CIGB), Cuba. The YFV Asibi strain and the WNV Israeli strain IS-98-STI were provided by the Biological resource Center of the Institut Pasteur. Stocks of DENV-2, YFV and WNV were produced on Vero NK cells. Viruses were concentrated by polyethylene glycol 6000 precipitation and purified by centrifugation in a discontinued gradient of sucrose. Flaviviruses were titrated on Vero NK cells by plaque assay as previously described [78] and titers were expressed in plaque-forming units (PFU)/ml. The Measles Schwarz strain expressing GFP (MeV-GFP) was described previously [40]. VSV Indiana and the CVB3 Nancy strain were kindly provided by N. Escriou (Institut Pasteur) and M. Bessaud, respectively (Institut Pasteur). Modified Vaccinia Ankara virus (MVA) expressing eGFP (MVA-GFP) was kindly provided by the ANRS via O. Schwartz (Institut Pasteur). It was manufactured by Transgene (Illkirch-Graffenstaden, France). The fluorescent marker, eGFP, is expressed under the control of the early promotor p11K7.5 and viral preparations were purified by tangential flow filtration. HMC3 cells were infected at the following MOIs: a MOI of 10 with DENV-2, 0,5 with WNV, 0,005 with VSV, 1 with MeV-GFP and 0,05 with MVA-GFP. IFN-treated HMC3 cells were infected at a MOI of 20 with DENV-2, 5 with WNV, 0,01 with VSV, 2 with MeV-GFP and 0,25 with MVA-GFP. Huh-7 were infected at a MOI of 1 with YFV and 0,25 with WNV. IFN-treated Huh-7 cells were infected at a MOI of 10 with YFV and WNV. Highly cell culture-adapted HCV Jad strain was obtained following transfection of Huh-7.5 cells with *in vitro* transcribed genome-length RNA as described previously [79–81]. Large volumes of HCV stocks were prepared following infection at a MOI of 0.01 50% tissue culture infectious doses 50 (TCID50) per cell with supernatants collected post-RNA transfection [82]. HCV infectious titers were determined by TCID50 assays in Huh-7.5 cells as described previously [79]. IFN-treated Huh-7.5 cells were infected at MOI of 3 TCID50/cell with HCV. The SARS-CoV-2 strain BetaCoV/France/IDF0372/2020 (historical) was supplied by the French National Reference Centre for Respiratory Viruses hosted by Institut Pasteur (Paris, France) and headed by Pr. S. van der Werf. The human samples from which the strain was isolated were provided by Dr. X. Lescure and Pr. Y. Yazdanpanah from the Bichat Hospital, Paris, France and Dr. Vincent Foissaud, HIA Percy, Clamart, France, respectively. A549-ACE2 cells were infected with SARS-CoV-2 at a MOI of 2.

### High throughput Screen

Five hundreds HMC3 cells were seeded in 384-well microplates in the morning of day 1 using a MultiDrop combi liquid dispenser (Thermo Fisher Scientific), in 40 μL of cell culture media. Cells were allowed to adhere for 4 hours (+/-1h) before transfection with individual siRNAs (10nM) diluted in a mix of OptiMEM (Life Technologies) and 0.05μL of Interferin reagent (Polyplus Transfection). siRNAs were transfected using an Evo 150 with MCA384 (Tecan). The library contained 1158 siRNA targeting 386 genes. siRNA targeting KIF11 was used to assess the transfection efficiency. On day 2, cells were treated with 1000U/ml of IFNα2a. Interferon was diluted into cell culture media and 10μL of the mix was robotically transferred to each well (except non-treated controls). 24 hours after IFN treatment, cell media was removed from siRNA-transfected plates and 40μL of the ZIKV PF13 strain, diluted to a final concentration of 7,500 particles/well, was added to the plates with the MCA384 head (Tecan). ZIKV titer was 6.5.10^8^ PFU/ml. Cells were then incubated 24 hours prior to fixation. Cells were fixed with 4% of formaldehyde (Sigma-Aldrich) for 20 min, plates were then washed once with PBS and quenched with NH4Cl (50mM) solution. Cells were then blocked with 1% BSA solution and permeabilized with 0.5% Triton X-100. Cells were next incubated for 60 min with mouse primary antibody anti-4G2 (1:500) which reacts with flavivirus E proteins. Cells were then washed twice in PBS solution and incubated with Alexa Fluor 488-coupled secondary antibodies (ThermoFisher Scientific). Nuclei were stained with 0.2 μg/ml Hoechst (Sigma). Images were acquired with an INCell2200 automated wide-field system (GE Healthcare,) using a Nikon 10X/0.45, Plan Apo, CFI/60. Three fields per well were analyzed using the INCell Analyzer 3.7 Workstation software. Two independent screens were performed. The mean cell count and the percentages of infected cells were extracted from quantification.

### Data analysis and hit calling

In the first analysis, data were processed using a software developed internally at the Biophenics platform. For hit identification, the robust Z-score method was used under the assumption that most siRNAs are inactive against ZIKV and can serve as controls [83, 84]. Raw values were log-transformed for cell count only to make the data more symmetric and close to normal distribution. In order to correct for plate positional effects, median polishing [84] was applied to each analyzed feature. It iteratively subtracts row, column and well median, computed from all plates within one screen. Hits for each compound were identified as follows: sample median and median absolute deviation (MAD) were calculated from the population of screening data points (named as sample) and used to compute Robust Z-scores (RZ-scores) according to a formula, in which the reference population corresponds to the siRNA-treated wells, and MAD is defined as the median of the absolute deviation from the median of the corresponding wells:

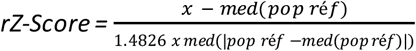

A gene was identified as a ‘hit’, if the RZ-score was < −2 or > 2 pointing in the same direction for 2 siRNAs targeting the same gene in both screens. Final values in the hit table correspond to the RZ-score of the second strongest siRNA. In the second analysis, data were process using an homemade script and CellProfiler [85]. Nucleus and viral assembly sites detected by the E signal were counted. As in the first analysis, rZ-Score and percentages of infected cells were quantified. Considering that each gene was targeted by three individual siRNA, genes were clusterized as hits, if at least two over three of their robust Z score absolute value were superior to 2. Genes were defined as hits when they were identified in at least one of the analysis.

### Antibodies, plasmids and reagents

The following primary antibodies were used in the study: anti-E MAb 4G2 hybridoma cells, anti-NS5-ZIKV [86], anti-VSV-G (IE9F9, Kerafast), anti-CVB3 VP1 (M7064, Agilent), anti-SARS-CoV-2 S protein mAb10 (1 μg/ml, a kind gift from H. Mouquet, Institut Pasteur, Paris, France), APOL1 (HPA018885, Sigma), MTA2 (8106, abcam), GM130 (12480, cell signaling), PI(4)P (Z-P004), PI(4)KB (06-578, Milipore) and anti-actin (A5316, Sigma). Secondary antibodies were as followed: anti-mouse Alexa 488 (A11001, Life Technologies), anti-mouse Alexa 680 (A21058, Life Technologies) and antirabbit DyLight 800 (SA5-35571, TermoScientific). The PI4KB inhibitor (1881233-39-1, MedChemExpress) and IFNα2a (Sigma-Aldrich, SRE0013) were used at the indicated concentration. GFP-APOL3 et APOL3-GFP were subcloned into pcDNA.3.1 from templates previously described [41].

### siRNA transfection

HMC3 cells were transfected using INTERFERin transfection reagent (Polyplus Transfection). Huh-7 cells, Huh-7.5 cells and podocytes were transfected with siRNAs at 10 nM final concentration using Lipofectamine RNAiMax (Life Technologies). All siRNAs were obtained from Dharmacon (siGENOME-SMARTpool).

### RNA extraction and RT-qPCR assays

Total RNAs were extracted from cell lysates using the NucleoSpin RNA II Kit (Macherey-Nagel) following the manufacturer’s protocol and were eluted in nuclease-free water. First-strand complementary DNA synthesis was performed with the RevertAid H Minus M-MuLV Reverse Transcriptase (Thermo Fisher Scientific). Quantitative real-time PCR was performed on a real-time PCR system (QuantStudio 6 Flex, Applied Biosystems) with SYBR Green PCR Master Mix (Life Technologies). Data were analyzed with the ΔΔCT method, with all samples normalized to GAPDH. All experiments were performed in technical triplicate. Viral genome equivalents concentrations (GE/ml) were determined by extrapolation from a standard curve generated from serial dilutions of the plasmid encoding the full-length genome of the Zika strain MR766 [87] or plasmids encoding a fragment of the RNA-dependent RNA polymerase (RdRp)-IP4 of SARS-CoV-2. HCV RNA was quantified by one-step reverse transcription-quantitative PCR using 50 ng of total intracellular RNA and TaqMan^®^ Fast Virus 1-Step Master Mix (Applied Biosystems) with primers and probe targeting the HCV 5’ nontranslated region as described previously [82]. Viral RNA levels were normalized with respect to 18S RNA levels quantified in parallel using TaqMan ribosomal RNA control reagents (Applied Biosystems). The products were analysed on a 7500 Fast Real-Time PCR system (Applied Biosystems). Serial dilutions of a genome-length *in vitro* transcribed HCV RNA served to establish standard curves and calculate HCV GE/μg total RNA concentrations. Primers and probe used for RT-qPCR analysis are given in Table S2.

### Western blot analysis

Cells were collected in RIPA buffer (Sigma) containing protease inhibitors (Roche Applied Science). Cell lysates were normalized for protein content with Pierce 660nm Protein Assay (Thermo Scientific), boiled in NuPAGE LDS sample buffer (Thermo Fisher Scientific) in non-reducing conditions. Samples were separated by SDS-PAGE (NuPAGE 4–12% Bis-Tris Gel, Life Technologies) with MOPS running buffer. Separated proteins were transferred to a nitrocellulose membrane (Bio-Rad). After blocking with PBS-Tween-20 0.1% (PBST) containing 5% milk for 1 h at RT, the membrane was incubated overnight at 4°C with primary antibodies diluted in blocking buffer. Finally, the membranes were incubated for 1 h at RT with secondary antibodies diluted in blocking buffer, washed, and scanned using an Odyssey CLx infrared imaging system (LI-COR Bioscience).

### Flow cytometry

Infected cells were fixed with cytofix/cytoperm kit (BD Pharmingen) and stained using the indicated primary and secondary antibodies. Non-infected, antibody-stained samples served as controls for signal background. Data were acquired using Attune NxT Acoustic Focusing Cytometer (Life Technologies) and analyzed using FlowJo software.

### Immunofluorescence assay

Cells were fixed with PFA 4% (Sigma) during 20min. Cells were permeabilized with PBS Triton X-100 (0.5%) for 15min at RT. After washing with PBS, they were incubated for 30 min with PBS + 0.05% Tween 20 + 5% BSA. The slides were then incubated overnight at 4°C with primary antibodies diluted in PBS. After washing with PBS, they were incubated for 1 h with secondary antibodies and washed with PBS. Nuclei were stained using PBS/NucBlue (Life Technologies, R37606). The mounting medium used is the Prolong gold (Life Technologies, P36930). All preparations were observed with a confocal microscope (ZEISS LSM 700 inverted) and images were acquired with the ZEN software.

### Statistical analysis

Data are presented as means ± SD and were analyzed using GraphPad Prism 7. Statistical analysis of percentage values or fold enrichment values were performed on logit or log-transformed values, respectively. Statistical analysis was performed with two tailed paired t-test or by one- or two-way analysis of variance (ANOVA) with Dunnet’s multiple comparisons test. Each experiment was performed at least twice, unless otherwise stated. Statistically significant differences are indicated as follows: *: p < 0.05, **: p < 0.01 and ***: p < 0.001; ns, not significant.

## Acknowledgments

We thank Dr. M.A. Saleem (University of Bristol, UK) for generously providing the podocytes (via E. Pays, Université Libre de Bruxelles, Belgium); C.M. Rice (Rockefeller University, New York, USA) for Huh-7.5 cells; Cinzia Traboni (IRBM, Pomezia, Italy) for Huh-7 cells; M. Bessaud (Institut Pasteur) for the CVB3 Nancy strain; N. Escriou (Institut Pasteur) for the VSV Indiana strain and anti-VSV-G antibodies; V. M. Cao-Lormeau and D. Musso (Institut Louis Malardé, Tahiti Island, French Polynesia) for the ZIKV-PF13 strain; L. Hermida and G. Enrique Guillen Nieto from the Centro de Ingeniería Genética y Biotecnología (CIGB), Cuba, for the DENV-2 strain Malaysia SB8553; T. Wakita (NIID, Tokyo, Japan) for pJFH1 HCV cDNA; R. Bartenschlager (University of Heidelberg, Germany) for pJFH1-2EI3-adapt cDNA; the French National Reference Centre for Respiratory Viruses hosted by Institut Pasteur (France) and headed by S. van der Werf for providing the historical SARS-CoV-2 viral strains; A. Merits (University of Tartu, Estonia) for anti-ZIKV NS5 antibodies; P. Desprès (Université de la Réunion, PIMIT) for 4G2 hybridoma cells; H. Mouquet (Institut Pasteur) for anti-SARS-CoV-2 S antibodies; M. Evans (Icahn School of Medicine at Mount Sinai, New York, USA) for the plasmid encoding the full-length Zika MR766 genome; F. Porrot and O. Schwartz (Institut Pasteur) for producing and sharing the MVA-GFP, C. Combredet (Institut Pasteur) for producing MeV-GFP; as well as E. Pays and S. Uzureau (Université Libre de Bruxelles, Belgium) for APOL1 and APOL3 plasmids and for stimulating APOL-focused discussions. We are grateful to the members of our laboratories for helpful discussions and technical advice. Finally, we thank Emeline Perthame (Bioinformatics and Biostatistics HUB, Institut Pasteur) for her help in statistical analysis.

## Declaration of interests

The authors declare no competing interests.

## Supplementary figure legends

**Figure S1. Quality control and reproducibility of the screens.** (A) HMC3 cells were transfected with either pool of 3 siRNAs against IFNAR or IFITM3 or non-targeting (NT) siRNAs, treated with IFNα2 (1000U/mL) for 24 hours and infected with ZIKV PF13 at a MOI of 7 PFU/cell for 24 hours. The number of cells positive for viral protein E was assessed by confocal analysis using the pan-flavivirus anti-E antibody 4G2. (B) Distribution of the “number of cells per 3 fields” parameter for each screen. The values of the control wells (cells transfected with siRNA targeting KIF11) are shown in dark gray. (C) Representation of the number of cells per 3 fields of screen 1 as a function of the screen 2. The green line represents the linear regression as compared to the expected perfect correlation (dotted black line). (D) Distribution of the “percentage of infected cells” parameter for each screen in the first analysis. The values of the control conditions (cells transfected with siRNAs targeting IFNAR1 or IFNAR2) are shown in dark grey. (E) Representation of the percentage of infected cells per well of screen 1 as a function of screen 2, as identified by the first analysis. The green line represents the linear regression as compared to the expected perfect correlation (dotted black line). (F) Representation of the percentage of infected cells per well in the analysis 1 as a function of analysis 2. The green line represents the linear regression. (G) HMC3 cells were transfected with pools of 3 siRNAs targeting the indicated genes or with non-targeting (NT) control siRNAs. The relative abundances of the mRNAs of the candidate genes were determined by RT-qPCR analysis and were normalized with respect to GAPDH mRNA level. They are expressed relatively to abundance in cells transfected with NT siRNAs set at 1. Data are means ± SD of three or four independent experiments. ND: not determined due to mRNA levels below assay threshold. The samples are the same than in Fig. 1E-H.

**Figure S2. Efficacy of the specific siRNAs.** Huh-7.5 cells (A) or A549-ACE2 cells (B) were transfected with pool of 3 siRNAs targeting the indicated genes or with non-targeting (NT) control siRNAs. The relative abundances of the mRNAs of the candidate genes were determined by RT-qPCR analysis and were normalized with respect to GAPDH mRNA level. Values are expressed relatively to abundance in cells transfected with NT siRNA in each experiment, set at 1. Data are means ± SD of three or four independent experiments. ND: not determined due to mRNA levels below assay threshold. The samples are the same than in Fig. 2.

**Figure S3. Localization of GFP-tagged version of APOL1 and APOL3 in HMC3 cells**. Cells were transfected with GFP-tagged versions of APOL1 and APOL3. Thirty hours later, they were stained with antibodies recognizing EEA1 or CD63 and with NucBlue to detect nuclei. Images are representative of numerous observations over 2 independent experiments.

## Notes

### Competing Interest Statement

The authors have declared that no competing interests exist.

