## Supplementary figures and images for "Discovery of genes that modulate flavivirus replication in an interferon-dependent manner"

### Supplemental figures

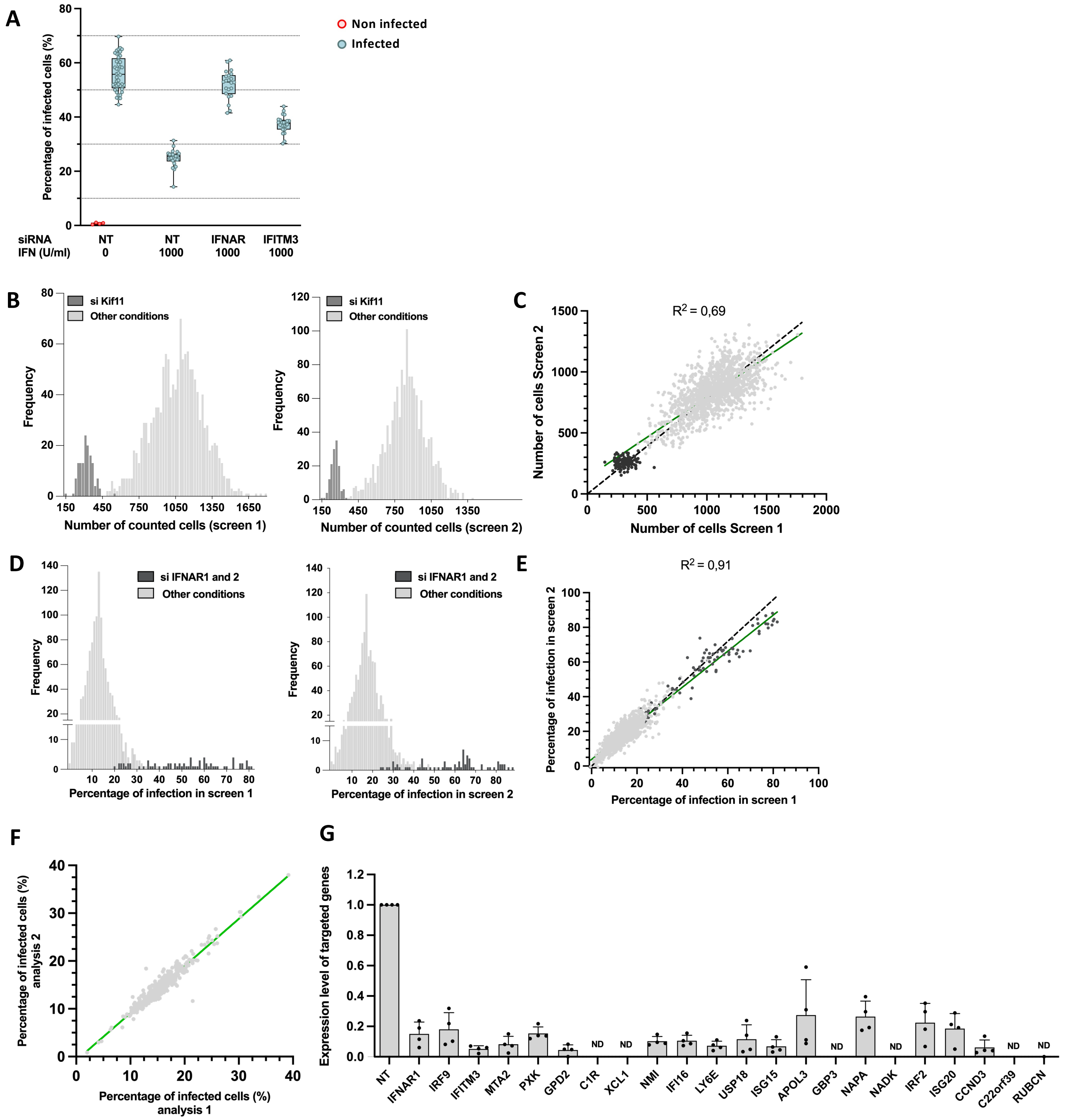

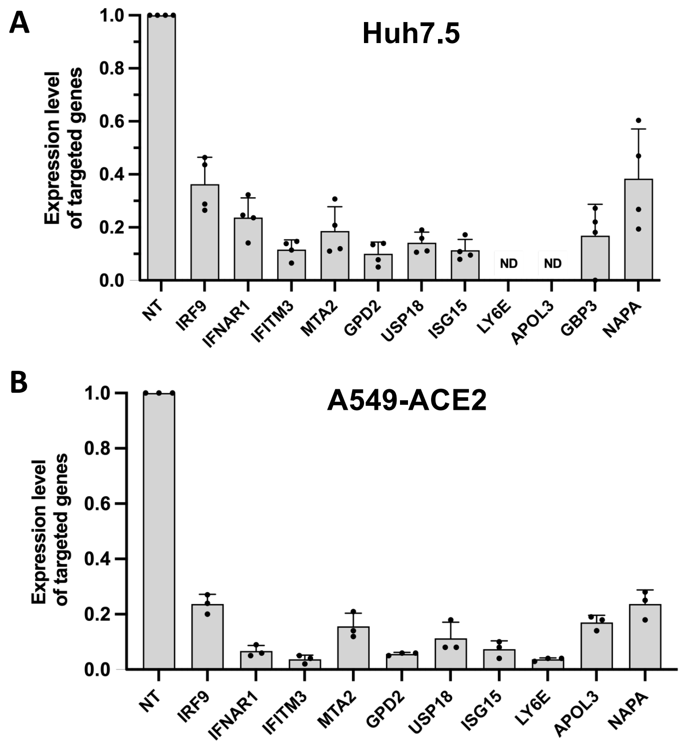

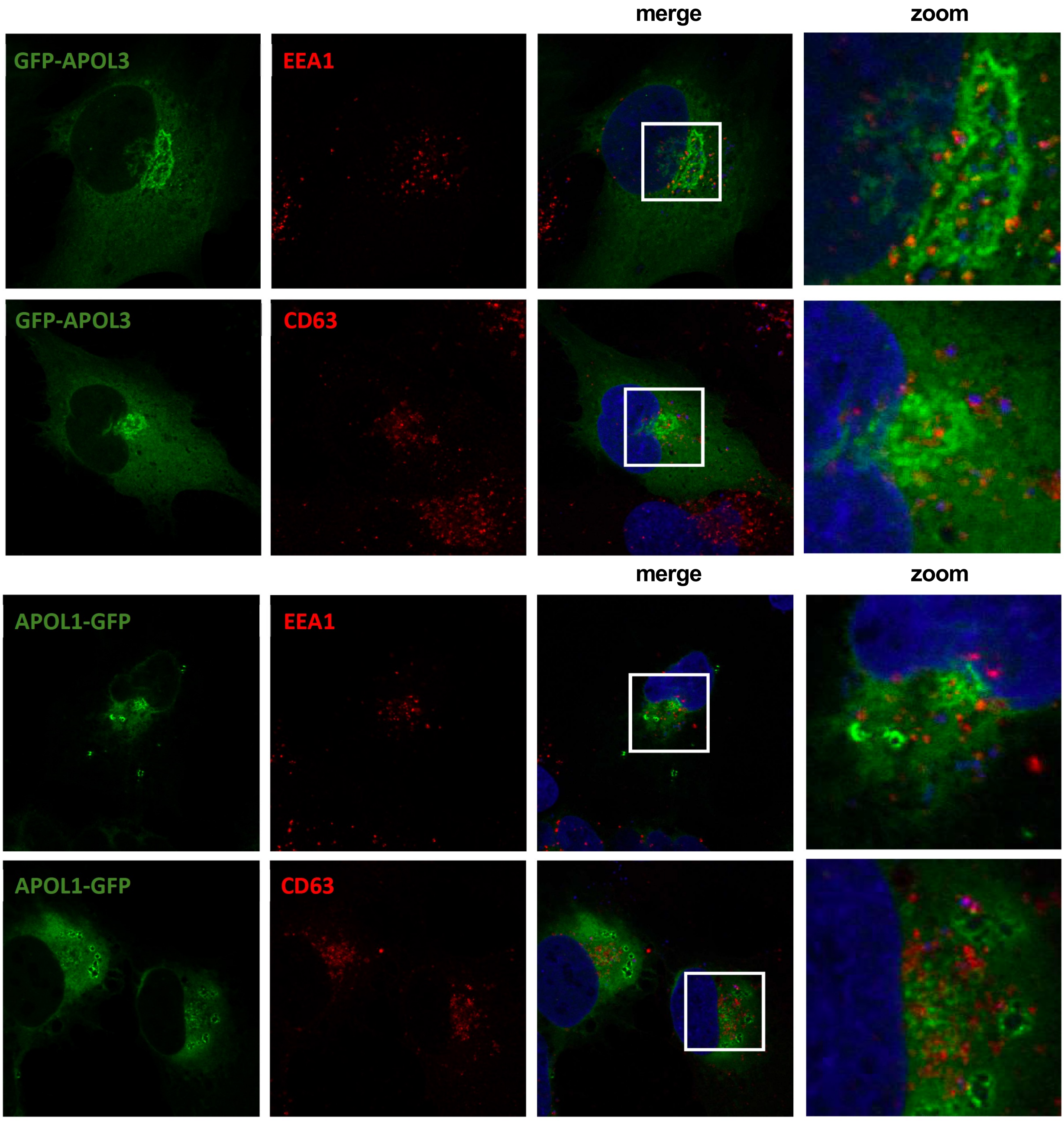
